# Functional Imaging of Developing Brain in Mice and Non-Human Primates

**DOI:** 10.1101/2024.06.07.596778

**Authors:** Jiejun Zhu, Dongming He, Mengzhu Sun, Hanming Zheng, Zihao Chen, Jin Yang, Chengqi Lin, Qiwen Yuan, Yun Stone Shi, Lei Sun, Zhihai Qiu

**Affiliations:** Guangdong Institute of Intelligence Science and Technology, Hengqin, Zhuhai, Guangdong 519031, China; Department of Biomedical Engineering, The Hong Kong Polytechnic University, Hung Hom, Hong Kong SAR, P. R. China, 999077; School of Life Science and Technology, Key Laboratory of Developmental Genes and Human Disease, Southeast University, Nanjing 210096, China; Obstetrics Department, The Fifth Affiliated Hospital of Sun Yet-Sen University, Zhuhai 519000, China

## Abstract

Despite significant advances in structural and genetic studies, investigations of early embryonic functions, such as brain activity, have long been constrained by technical challenges. Functional ultrasound (fUS) has emerged as a breakthrough modality, enabling real-time monitoring of brain activity with exceptional spatial and temporal resolution and offering unprecedented opportunities for studying functional embryonic development. In this study, we used fUS to monitor whole-embryo activity in mice from embryonic days E8.5 to E18.5, revealing patterns of neural activity throughout embryogenesis. This approach provides new opportunities to explore brain development dynamically as it unfolds. Moreover, we observed embryo responses to external stimuli, including sound, in both mice and cynomolgus macaques, offering insights into early sensory processing and neural maturation. In summary, our study establishes fUS as a powerful tool for studying embryonic brain functional development, with significant implications for scientific research, especially in non-human primate models, and clinical applications.

## Introduction

Much of the current understanding of mammalian embryonic development has been gained from studying animal anatomical dissections and organoids^1–3^, which have revealed critical time-points for embryogenesis, organogenesis, and their corresponding gene factors. For example, the auditory system—including sensory transduction in hair cells^4^, the auditory nerve in the peripheral system, and central nervous system connections^5,6^—has been extensively studied and shown to be partially developed before birth. However, its *in-vivo* activity remains less well understood. Metabolic states, similarly, have been found to regulate vascular and neural development at different embryonic stages^7–9^ though this process has never been directly observed *in vivo*. Modern molecular biological methods such as single-cell RNA sequencing and spatial transcriptomics have further enriched our understanding, including uncovering the neuronal-type hierarchy and the potential selective vulnerability for neurodevelopmental disorders, including autism^10^. However, these developmental disorders are limited by discontinuous temporal observations and the inability to track changes in real time.

To address this problem, several kinds of functional imaging technologies have been developed to gain deeper insight into fetal development. Optical methods, including two-photon microscopy and light-sheet microscopy, have proven invaluable for imaging *C. elegans*, *Drosophila* and zebrafish^11^, providing detailed insights into whole-embryo cellular-level development. However, when it comes to applying optical methods to mammalian embryos, the ability to gain a comprehensive view of the whole embryo becomes curtailed^12^. Recently, a study utilized photoacoustic microscopy to capture both the structure and function of the heart during mouse gestational days (GDs) 14.5–17.5^13^, while another study using the same method focused on capturing placental function under both normal and pathological conditions^14^. But, these methods are limited in the transparency they offer and, thus, do not permit whole-embryo observation. Without a holistic perspective, how different organs function, interact, and coordinate with each other during development, ultimately culminating in the successful emergence of a new life cannot fully be grasped. Magnetic resonance imaging (MRI) provides sufficient coverage of the entire fetus^15^ but primarily evaluates only its structural features. A modified approach, functional magnetic resonance imaging (fMRI), is capable of exploring the functional characteristics of the brain^16–18^, but it has limited application in the fetal context due to insufficient resolution and motion artifacts^19,20^. Nevertheless, the fundamental principle of fMRI remains sound - it assesses hemodynamic changes that mirror brain function, operating under the assumption that the heightened energy demand during brain activity necessitates circulatory support^16^.

Building upon this principle, fUS was recently introduced as a modality for real-time monitoring brain activity in rodents and macaques^21–23^. fUS offers a distinctive blend of attributes, including expansive spatial coverage (up to 10 cm), high spatiotemporal resolution (down to 100 μm and up to 10 ms), and ample sensitivity (detecting relative hemodynamic changes of as little as 2% without the need for averaging across multiple trials)^22–24^. Thus, fUS as a modality offers the potential for effective, comprehensive monitoring of fetal hemodynamic activity with high spatiotemporal resolution without being hampered by typical anatomical limitations. High-frequency ultrasound (HFUS) has also been widely utilized for imaging embryonic development in mice, providing detailed insights into cardiovascular function and organogenesis, particularly from embryonic day 8-8.5 (E8-E8.5) onwards^25–27^. Studies by Phoon et al. and others have demonstrated the utility of HFUS in visualizing the mouse heart and vasculature, capturing important developmental milestones with spatial resolutions down to 30 μm^28–30^. However, while HFUS offers high resolution, its application is primarily limited to structural and flow imaging, lacking the functional insights provided by newer modalities like fUS.

In this study, we aimed to establish fUS as a tool to monitor functional activities of developing mouse and cynomolgus macaque embryos. Our data revealed that fUS effectively images the entire mouse embryo, offering exceptional sensitivity and spatiotemporal resolution from E8.5 to E18.5. By deconvoluting brain fUS signals from those of the heart, we were able to capture independent brain activities across the gestational period, with an increase of delta rhythm observed from E13.5 and a slight decrease in E18.5. More importantly, fUS recordings revealed that the developing mouse brain exhibits responsiveness to various external stimuli, including auditory cues. Notably, our study identified sound-induced brain activity as early as embryonic day 18.5 (E18.5) in mouse embryos, challenging traditional knowledge. Furthermore, by utilizing fUS imaging in cynomolgus macaque embryos, we detected brain responses to auditory stimuli within the auditory pathway. This analysis provided critical insights into the early development of the auditory pathway and motor system. Taken together, our study provides novel insights into embryonic brain functional development. With the existent widespread use of fetal ultrasound in clinical settings, our findings lay a foundation for advancing fetal examination techniques and possible early detection of disorders or conditions.

## Results

### fUS Imaging Captures Embryonic Functioning with Exceptional Spatial and Temporal Resolution Across Depth

fUS utilizes plane-wave ultrasound to generate accurate, high-resolution images of mouse embryos in vivo in a fully non-invasive manner (Movie S1). To simulate in utero ultrasound imaging as performed in clinical settings^31,32^, this study employed a minimally invasive approach (Fig. 1). By acquiring 25 compound images at 500 Hz, each derived from seven planar illuminations tilted from −6° to 6° using a 15-MHz ultrasound probe, we successfully imaged the entire mouse embryo (128 × 80 mm) at an impressive imaging rate of 20 Hz. This technique enabled a time-resolved readout of blood volume variations within each voxel across the entire embryo, providing a detailed and dynamic view of embryonic physiology.

**Fig. 1:**
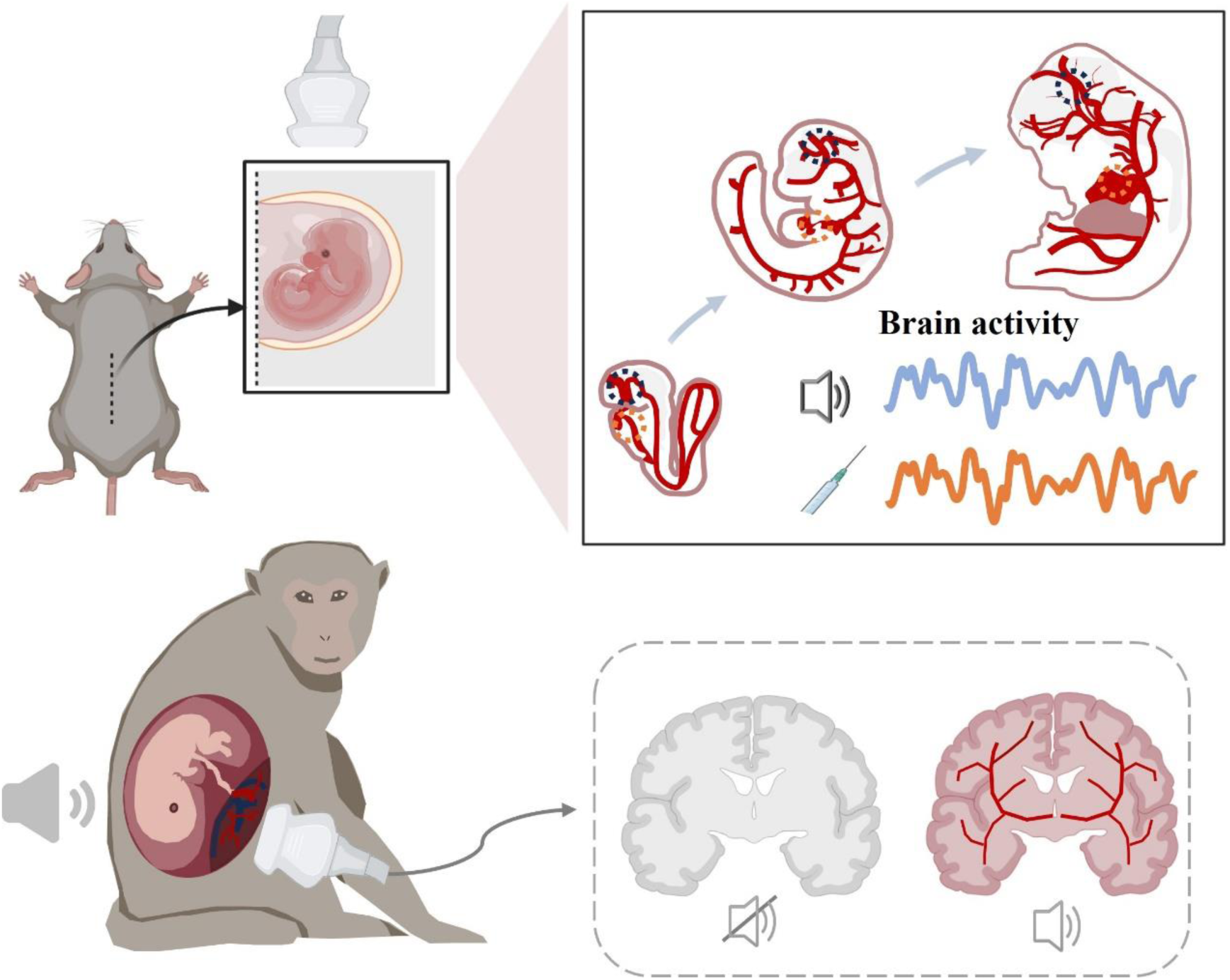
Schematic representation of embryonic functional ultrasound (fUS) imaging throughout the gestational period. Schematic representation of a pregnant mouse (top left) and a cynomolgus macaque (bottom left) undergoing fUS imaging. A mouse embryo, still inside the uterus, was extracted through a small incision in the pregnant mouse’s abdomen (middle left). Embryos ranging from E8.5 to E18.5 were imaged either under resting conditions or in response to different external stimuli, and their cardiac function and brain activity were analyzed (right). The brain activity of three cynomolgus macaque at different gestational days (30, 85, and 85) was imaged with auditory stimulation.

Compared to traditional line-by-line B-mode ultrasound imaging, fUS imaging provides significantly richer information about the embryonic circulatory system by analyzing 2-second fUS signals, comprising 1,000 compound images (Fig. 2A-D). Furthermore, the use of stepped scanning allows for the clear visualization of the embryonic circulatory system in sagittal, coronal, and horizontal planes, enabling detailed functional observations in regions such as the placenta, spinal cord, and brain vascular networks (Movies S2, S3, and S4).

**Fig. 2:**
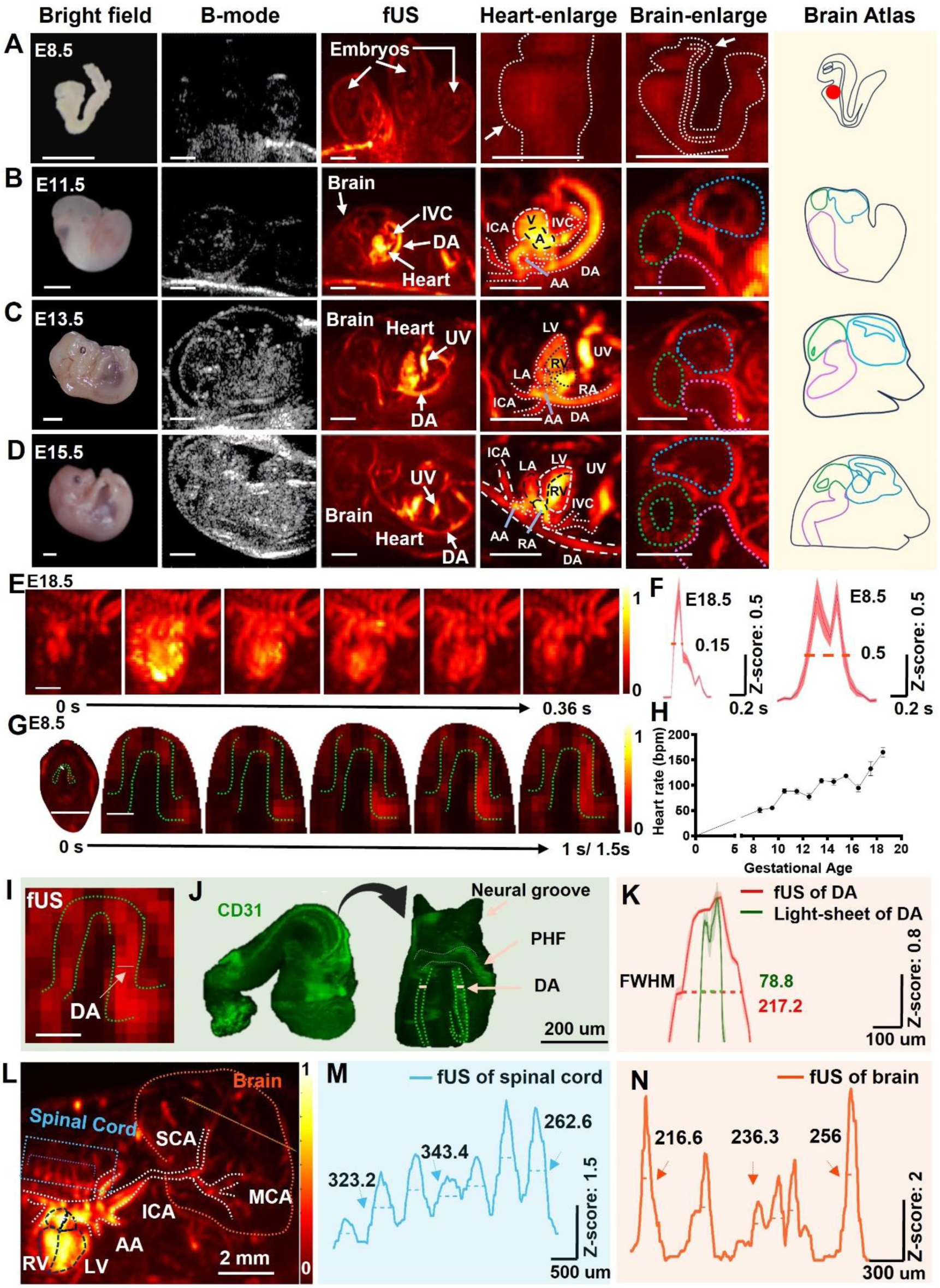
Spatial-temporal fUS imaging of developing embryonic mouse heart and brain. A) Representative images of E8.5 mouse embryos under (from left to right) bright field, B-mode ultrasound, fUS, heart fUS enlarged, brain fUS enlarged, and the corresponding brain atlas images. Scale bars: 2mm. B) Representative images of E11.5 mouse embryos. Scale bars: 2mm. C) Representative images of E13.5 mouse embryos. Scale bars: 2mm. D) Representative images of E15.5 mouse embryos. Scale bars: 2mm. E) Representative image sequence of heart fUS signals within a heartbeat cycle obtained from an E18.5 embryo. Scale bar: 1mm. F) Averaged fUS signal of the heart within a heartbeat cycle in E18.5 mouse embryo, which FWHM is 0.15 seconds (left). Average fUS signal of the heart within a heartbeat circle in a E8.5 mouse embryo, which FWHM is 0.5 seconds. G) Representative image sequence of heart fUS signals within a heartbeat cycle obtained from an E8.5 embryo. Scale bar: 2mm & 1mm. H) Summary of embryonic heartbeats ranging from E8.5 to E18.5 (n=3-4 in each group). Data presented as Mean ± SEM. I) Representative imaging frame of fUS data obtained from an E8.5 embryo within a heartbeat cycle. Green dotted lines delineate the dorsal aorta (DA). White dotted line indicates the DA region (DA diameter) targeted for further analysis. Scale bar: 1mm. J) Light-sheet imaging of an E8.5 embryo stained with CD31 (an endothelial marker of blood vessels). Green dotted lines delineate the embryonic DA. White dotted lines indicate the DA region (DA diameter) targeted for further analysis. K) Signal intensity along the vertical axis of the dorsal aorta of the E8.5 embryo presented in I and J panel. FWHM of fUS is 217.2 μm, and the FWHM of light-sheet imaging is 78.8 μm. L) Representative imaging frame of fUS data obtained from an E15.5 embryo. White lines dotted lines indicate major organs and vessels. Blue dotted line and orange dotted line indicate the spinal cord and a randomly drawn line across the brain vessels for subsequent analysis. M) Distribution of fUS signal intensity of spinal cord blood flow along the blue lines in panel L. The FWHM values are 242.42, 323.23, 242.42, 343.44, 282.82, 242.43 and 262.62 μm. N) Distribution of fUS signal intensity of brain blood flow along the origin lines in panel L. The FWHM values are 216.61, 216.61, 236.31, 295.39, 236.3, and 256 μm. **Abbreviations in** Fig. 2: A, atrium; AA, aortic arch; DA, dorsal aorta; ICA, internal carotid artery; IVC, inferior vena cava; LA, left atrium; LV, left ventricle; MCA, middle cerebral artery; PHF, primary heart field RA, right atrium; RV, right ventricle; SCA, superior cerebellar artery; UV, umbilical vein; V, ventricle.

Using fUS, we visualized cardiac function in E18.5 embryos with a heartbeat cycle of approximately 0.36 seconds (Fig. 2E). The full-width at half maximum (FWHM) of the heart’s fUS signal was measured at 0.15 seconds (Fig. 2F, left), highlighting its high spatiotemporal resolution. Notably, fUS captured embryonic circulation as early as E8.5 (Fig. 2G), when the heart is still in the tubular stage (Movie S5). Individual heartbeats were accurately recorded, with an FWHM of 0.5 seconds (Fig. 2F, right).

Analysis of fUS signal intensity in the dorsal aorta revealed a FWHM approximately 2.76 times greater than that of the corresponding immunofluorescence (IF) signal from CD31 vascular staining (217.2 µm vs. 78.8 µm) (Fig. 2I-K). This superior precision provides sufficient resolution to investigate functional changes in microcirculation systems throughout the gestation period. Additionally, fUS successfully visualized vessels in the mouse spinal cord and brain (Fig. 2L), with FWHM measurements of 277.05 ± 15.72 µm (Mean ± SEM) in the spinal cord (Fig. 2M) and 242.87 ± 27.07 µm (Mean ± SEM) in the brain (Fig. 2N). These results underscore the exceptional spatial resolution of fUS, enabling the detection of functional dynamics in embryonic heart and brain systems.

### Embryonic Brain Activity Can be Monitored Through fUS Imaging Independent of Heart Pulses

Using B-mode ultrasound to navigate embryo positioning and analyze the features of major cerebral arteries (such as the anterior cerebral artery and posterior cerebral artery), alongside fUS imaging of the brain anatomy, we successfully segmented the brain into forebrain, midbrain, and hindbrain regions (Fig. 3A). Remarkably, fUS enabled simultaneous study of both heart and brain signals within a single embryo (Fig. 3B). Our analysis revealed a strong influence of cardiac activity on brain signals (Fig. 3C-E), with each heartbeat cycle coinciding with brain activity (Movie S6). The correlation between heart and hindbrain signals reached a maximum of 0.78 at E12.5 (Fig. 3D-E).

**Fig. 3:**
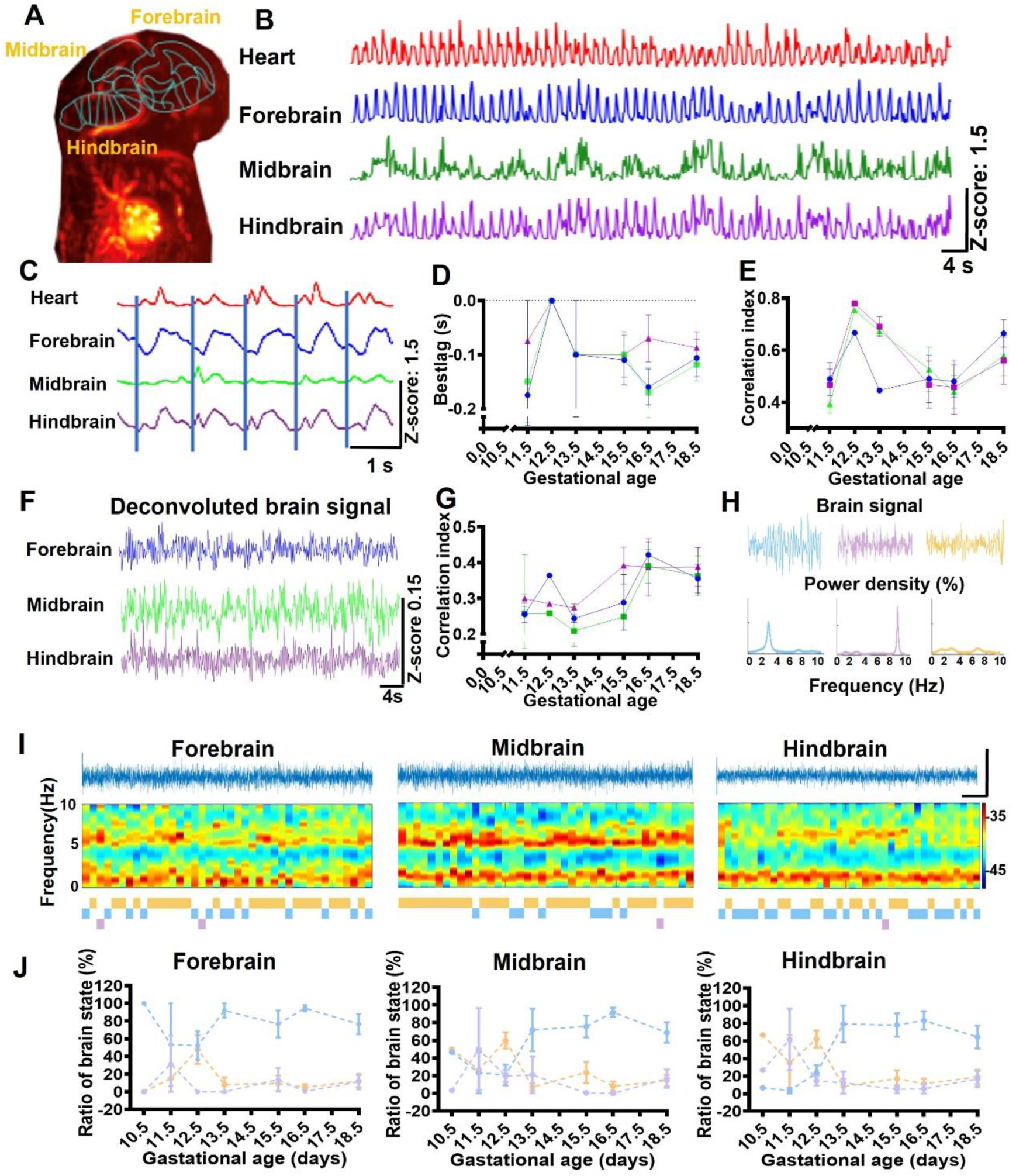
Deconvolution analysis of fUS signals reveals central nervous system activities during mouse embryogenesis. A) Representative brain atlas image of E15.5 mouse embryos obtained through fUS. B) Hemodynamic changes are presented as Z-scores. From top to bottom, traces represent heart (red), forebrain (blue), midbrain (green), and hindbrain (purple). C) Exemplars of fUS signals showing a high correlation between the heart and forebrain, midbrain, and hindbrain (from top to bottom). The brain fUS signals follow the onset of heart activity. D) Delay in forebrain (blue), midbrain (green), and hindbrain (purple) activity relative to the heart during E11.5 to E18.5 embryonic days (n=2-8 in each group). Data presented as Mean ± SEM. E) Summarized correlation indices between the heart and forebrain (blue), heart and midbrain (green), and heart and hindbrain (purple), ranging from E11.5 to E 18.5 (n=2-8 in each group). Data presented as Mean ± SEM. F) Exemplars of brain signals in the forebrain (blue), midbrain (green), and hindbrain (purple) (from top to bottom) elucidated after deconvolution analysis. G) Correlation indices between the three brain regions, i.e., forebrain vs midbrain (blue), midbrain vs hindbrain (green), hindbrain vs forebrain (purple) are summarized ranging from E11.5 to E 18.5 (n=1-8 in each group). Data presented as Mean ± SEM. H) Principal classification of recorded brain states. Blue signals represent typical slow waves exhibiting a delta rhythm below 4 Hz, purple signals represent typical rapid waves within the theta range of 6 to 10 Hz, and yellow signals represent other epochs with a more even rhythm distribution. I) Representative signal traces, frequency spectrum and brain state classified every 10 seconds of forebrain, midbrain, and hindbrain (left to right) brain states. Scale bar: 40s & Z-score=05. J) Summary of forebrain, midbrain, and hindbrain activity (left to right) during the gestational period. Blue: delta rhythm wave; purple: theta wave; yellow: even rhythm. Data presented as Mean ± SEM.

While previous studies have demonstrated that fUS reflects neuronal activity in adult mouse brains^33^, our findings indicate that cardiac signals have a pronounced impact on fUS signals in developing mouse embryos. To mitigate artifactual signals induced by blood flow pulsation, we performed a deconvolution analysis. Post-deconvolution, distinct brain signals were discernible across all brain regions (Fig. 3F). Before deconvolution, brain responses closely followed heartbeats within a 0–200 ms window, as indicated by the optimal lag for achieving the highest correlation (Fig. 3D-E). After deconvolution, cross-correlation analysis among different brain regions (e.g., forebrain to midbrain, forebrain to hindbrain, and midbrain to hindbrain) revealed similar correlation patterns throughout gestation, with the lowest levels at E13.5 and the highest at E16.5 (Fig. 3G).

We further classified brain signals into three distinct activity patterns: slow waves exhibiting delta rhythms below 4 Hz, rapid waves within the theta range of 6–10 Hz, and other epochs with evenly distributed rhythms (Fig. 3H). Temporal analysis of these patterns revealed fluctuations in brain activity throughout development, with a significant prevalence of delta rhythms emerging at E13.5 and slightly decreasing by E18.5 (Fig. 3I-J). Notably, these changes correspond to known metabolic shifts in the brain during this developmental period^7–9^.

fUS, with its ability to monitor whole-embryonic functional activity during development, has the potential to act as an interface between the embryo in utero and the external world. To explore this concept, we encoded brain activity from different regions into three audible sound frequencies (Mm. 1). This approach allows humans outside to intuitively interpret brain activity through sound as a readout of embryonic brain states, offering a novel and potentially practical method for mothers to monitor fetal brain activity.

### fUS Detection of Brain Activation by External Stimuli in Developing Mouse Embryos

We further examined the functional state of the embryonic brain beyond resting-state analysis, aiming to investigate sensory responses to external stimuli. The study began with the administration of alcohol (3 g/kg) to pregnant mice, which induced marked changes in embryonic brain activity that were easily detectable without requiring deconvolution analysis (Fig. S1). To confirm the embryonic brain’s response to external stimuli, we utilized wide-field calcium imaging, a widely used method, and tested its response to the same sound stimulations. Sound stimuli were delivered to E18.5 embryos (Fig. 4A). Using fUS, we recorded brain activity in the sagittal view, clearly visualizing the forebrain, midbrain, and hindbrain. In parallel, wide-field calcium imaging captured activity in the cortex and midbrain. The midbrain, containing the Inferior Colliculus (IC), a critical component of the auditory pathway, was selected as the region of interest for further analysis, as this region was captured by both methods (Fig. 4B).

**Figure 4:**
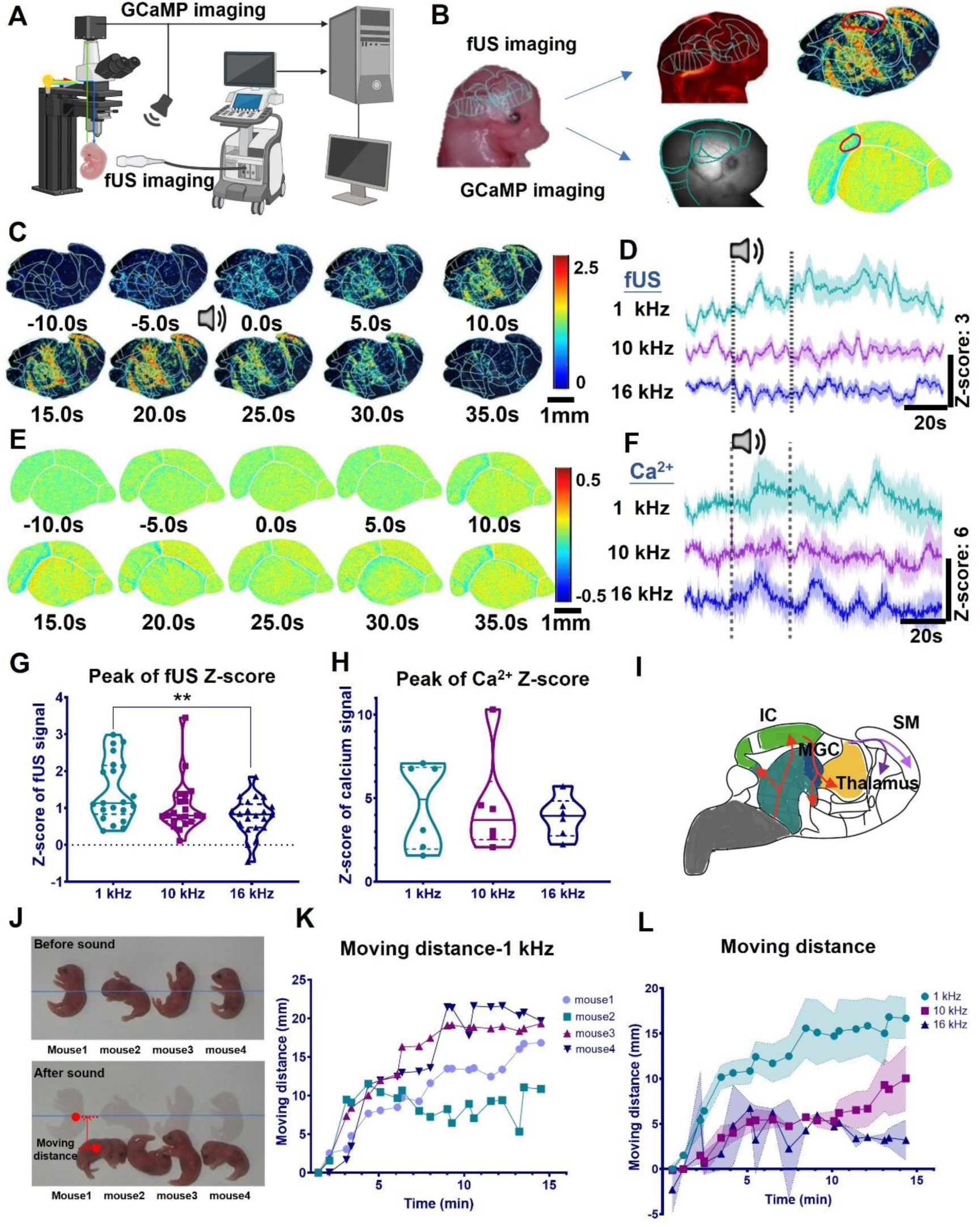
fUS detection of brain activation by external stimuli in developing mouse embryos. A) Schematic of simultaneous fUS and wide-field calcium imaging of E18.5 mouse embryo. B) Schematic of a mouse undergo fUS and wide-field calcium imaging (left). Representative images of embryonic mouse brain captured by fUS (top right) and by wide-field calcium imaging (bottom right). C) Representative fUS images showing the embryo brain’s response to a 25-second auditory stimulation (1 kHz, 70 dB SPL). Scale bar: 1mm. D) Z-score trace of the fUS signal in the midbrain during three 25-second auditory stimulations at 1 kHz, 10 kHz, and 16 kHz, from top to bottom (n=7 from 2pregnant mice). Data presented as Mean ± SEM. E) Representative calcium imaging images showing the embryo brain’s response to a 25-second auditory stimulation (1 kHz, 70 dB SPL). Scale bar: 1mm. F) Z-score trace of the calcium imaging signal in the midbrain during three 25-second auditory stimulations at 1 kHz, 10 kHz, and 16 kHz, from top to bottom (n=7 from 2pregnant mouse). Data presented as Mean ± SEM G) Summary of peak Z-scores for the fUS signal within 1 minute after the start of a 25-second auditory stimulation (n=7 from 2 pregnant mice). Data presented as Mean ± SEM. H) Summary of peak Z-scores for the calcium imaging signal within 1 minute after the start of a 25-second auditory stimulation (n=7 from 2 pregnant mice). Data presented as Mean ± SEM. I) Diagram of brain regions activated by auditory stimulation, including the IC, MGC, SM, and thalamus. J) Representative image showing movement of four P1 mice before and after sound stimulation (9 trials of auditory stimulation. Each trial includes a 25-second stimulation with a 1-minute interval, at 1kHz and 90 dB SPL). K) Summary of movement distance for four P1 mice in response to auditory stimulation at 1 kHz, 90 dB SPL. L) Summary of movement distances for P1 mice in response to auditory stimulation at 1 kHz, 10 kHz, and 16 kHz.

Figure 4C presents a series of representative fUS images showing the embryo brain’s response to 1 kHz sound stimulation. These sagittal views of neural activity revealed significant activation in key auditory pathway components, including the Inferior Colliculus (IC), Medial Geniculate Complex (MGC), and Thalamus. When additional auditory stimuli (1 kHz, 10 kHz, and 16 kHz) were applied, embryos responded to all frequencies, with distinct temporal dynamics in brain activity depicted as increases in Z-scores (Fig. 4D). Similarly, Figure 4E shows the corresponding calcium imaging series for the embryo brain’s response to 1 kHz sound stimulation. Responses to 10 kHz and 16 kHz stimuli were also observed, with consistent activity patterns across all frequencies (Fig. 4F). Notably, Z-score peaks from calcium imaging demonstrated patterns comparable to those observed with fUS (Fig. 4G-H). Analysis of sound-induced brain responses revealed heightened sensitivity to 1 kHz stimuli compared to 16 kHz stimuli (P = 0.006) (Fig. 4G). These findings provide direct evidence of embryo brain sensory responses to external auditory stimulation.

Interestingly, both fUS and calcium imaging revealed neural activity extending to the somatomotor (SM) area (Fig. 4I), suggesting broader involvement of both auditory and motor circuits. This led us to hypothesize that auditory stimulation might trigger movement in neonatal mice. To test this, we recorded P1 mouse movements in response to 1 kHz, 10 kHz, and 16 kHz sound stimuli. Indeed, we observed movement away from the trumpet source of the sound (Fig. 4J). Repeated stimulations demonstrated that mice consistently moved farther from the sound source with increasing stimulation (Fig. 4K). In alignment with our fUS and calcium imaging data from E18.5 embryos, neonatal mice exhibited robust responses to sound stimuli, particularly to 1 kHz sounds (Fig. 4L). These results challenge previous findings suggesting that mice can only hear after P7/P8^34^.

In summary, these results demonstrate that fUS can reliably capture real-time neural responses, offering novel insights into auditory sensory processing and highlighting its potential for mapping functional connectivity in the fetal brain.

### fUS Detection of Sound-induced Brain Activities in Fetal Cynomolgus Macaques

To further explore the clinical potential of fUS, we performed functional imaging on three pregnant cynomolgus macaques: one at 30 days of gestation and two at 85 days. Traditional ultrasound (3.5 MHz), commonly used in clinical practice, successfully captured structural and Doppler images, clearly visualizing embryo structures. However, it provided limited brain Doppler signals in 85-day-old macaques (Fig. 5A, left). In contrast, fUS (3.5 MHz) allowed the recording of both cardiac activity and brain signals, albeit with lower spatial resolution (Fig. 5A, right). To achieve higher spatial resolution, we employed a robotic arm-controlled fUS (7.5 MHz) (Fig. 5B), successfully imaging blood flow across the brain vasculature (Fig. 5C-D). This confirmed the ability of fUS to detect detailed brain activity, though asymmetry in the right side was noted due to the imaging plane not being strictly perpendicular.

**Figure 5:**
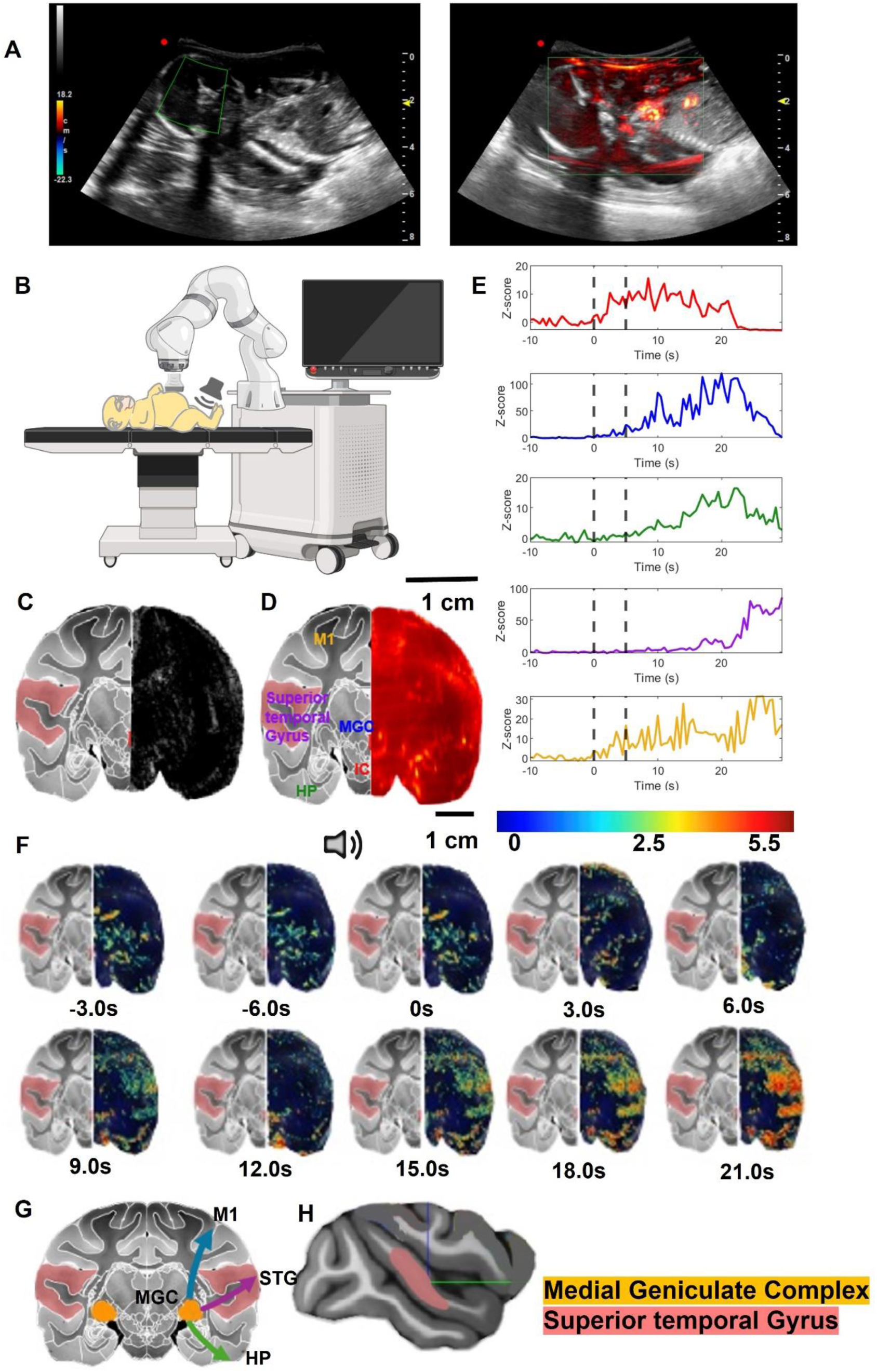
fUS detection of sound-induced brain activities in fetal cynomolgus macaque. A) Representative image of an 85-day-old embryonic cynomolgus macaque brain under traditional B-mode and color Doppler ultrasound (probe center frequency: 3.5 MHz) (left). Representative image of the embryonic cynomolgus macaque brain under fUS imaging with circular wave (probe center frequency: 3.5 MHz) (right). B) Schematic of an cynomolgus macaques undergo functional imaging with a 5-second 1kHz auditory stimulation. C) Representative image of B-mode (right hemisphere) (probe center frequency 7.5 MHz) and its corresponding brain atlas. Scale bar: 1cm. D) Representative image of the embryonic cynomolgus macaque brain under fUS imaging (right hemisphere) (probe center frequency 7.5 MHz) and its corresponding brain atlas. E) Z-score trace of fUS signal within a 30 second period with a sound stimulus at 10 seconds. F) Representative image sequence of brain function responses to a 5-second auditory stimulation. Scale bar: 1cm. G) STG location in a cynomolgus macaque brain (side view). H) Diagram of brain function responses to auditory stimulation, starting from IC, MGC and propagate to STG, hippocampus and motor cortex.

For the macaque at 30 days of gestation, traditional ultrasound (3.5 MHz) effectively navigated the embryo and identified umbilical cord structures, while fUS (7.5 MHz) successfully measured umbilical cord blood flow and detected umbilical cord entanglement (Fig. S2). However, due to the small size of the brain at this stage, fUS at 7.5 MHz was insufficient to visualize brain activity, leading us to focus on the 85-day gestation period.

We further analyzed temporal changes in fUS signals across the brain in response to external auditory stimuli. Following sound stimulation, we observed a strong initial response in the Inferior Colliculus (IC), which then propagated across the brain (Fig. 5E). The primary pathway involved the IC-Medial Geniculate Complex (MGC)-Superior Temporal Gyrus (STG), traversing key auditory nuclei (Fig. 5F-G). In addition, responses extended to the hippocampus and motor cortex, indicating higher-order brain functions. A schematic of auditory-induced brain activity is shown in Fig. 5G-H, providing clear evidence that both mouse embryos (E18.5) and cynomolgus macaque embryos (85 days) exhibit functional responses to external stimuli.

These findings highlight the early development of sensory processing pathways in the fetal brain, demonstrating the potential of fUS to explore brain function in utero. This capability is particularly relevant for understanding the early development of consciousness and higher-order brain functions.

## Discussion

The fetal stage is increasingly recognized as a critical period in the development of numerous diseases, such as autism^35,36^. However, monitoring fetal development, particularly brain development, remains a significant challenge, and effective treatment options are limited. Our study introduces fUS imaging, a novel technique that builds on the principles of clinical fetal ultrasound while providing unprecedented functional insights into brain activity throughout gestation. This represents a major advancement in embryonic imaging, with the potential to transform fetal examination practices in clinical settings.

Using fUS, we successfully captured the resting state of embryonic brain activity. By incorporating heartbeat signal deconvolution, we tracked resting-state brain activity across gestation and identified the emergence of a delta rhythm at E13.5, which slightly declined by E18.5. This rhythm corresponds to the NREM sleep patterns observed in adult EEG^37^ and aligns with predictions from data on premature infants and adults^38^, suggesting increased brain activity during late embryonic stages. The emergence of the delta rhythm coincides with key developmental milestones^7–9^, including vascularization in the ventricular zone, the transition of cortical neural progenitor cells (NPCs) to neurogenic divisions, and their metabolic shift to aerobic respiration at E13.5. By E16.5, some radial glial progenitors (RGPs) begin gliogenesis. These findings suggest that the delta rhythm may reflect these processes and serve as a marker for evaluating embryonic brain development.

Our study also demonstrated significant brain activity in response to external stimuli, such as alcohol and sound, without the need for deconvolution. fUS signals closely correlated with calcium imaging data, underscoring its reliability as an alternative method for measuring brain activity during embryonic development. Notably, we found that E18.5 mice respond to sound stimulation, as evidenced by fUS, calcium imaging, and behavioral assessments. This finding challenges previous auditory brainstem response (ABR) studies, which suggested that hearing only emerges after birth after (∼P7/P8)^34^. The auditory pathway and mechanotransduction in inner hair cells are already developed by E17^4–6^. The discrepancy with ABR results likely arises from methodological differences: ABR detects synchronized neuronal activity, whereas fUS resolves single-trial, phase-independent responses. fUS thus offers a powerful tool to study embryonic brain activity, providing unique insights into early brain function and sensory processing.

In exploring the clinical potential of fUS, we extended its application to non-human primates and successfully recorded auditory pathway responses to sound stimulation. This success further supports the feasibility of fUS imaging for clinical translation. Disorders such as autism are believed to originate during the embryonic stage, yet in-vivo detection methods remain unavailable. By leveraging fUS, we may be able to identify specific brain regions and critical timepoints for the onset of such disorders.

By capturing brain activity in both resting states and in response to external stimulation across mice and non-human primates, our study paves the way for early detection of fetal diseases, developmental disorders, and other conditions. The dynamic functional changes recorded with fUS could facilitate the development of embryo-computer interfaces and non-invasive interventions, such as ultrasound neuromodulation. These advancements promise to deepen our understanding of embryonic development and developmental disorders, potentially enabling earlier intervention and improved outcomes.

## Conclusion

In our study, we discovered that fUS effectively captures the functioning of the entire embryo throughout the mouse gestation period, offering exceptional sensitivity and spatiotemporal resolution. Independent of signals from heart pulsations, we were able to capture intricate and specific patterns of brain activity, both in resting state as well as under different stimuli. Specifically, we found that mouse fetuses respond to sound stimulations, providing new insight of sensory development. In addition, when fUS imaging was applied to cynomolgus macaques, the propagation of brain activity within the auditory pathway could be recorded. The successful capture of brain activity opens new avenues for research into embryonic brain activity, providing valuable insights into patterns of early brain development. In summary, our study demonstrates fUS as a useful tool for investigating embryonic functional development, paving the way for further studies of fetal development in primates, including humans.

## Supporting information

3D images of a whole embryonic mouse functioning

Functional imaging of an E15.5 mouse

Functional imaging of an E8.5 mouse

## Author contributions\

Conceptualization: ZHQ, JJZ, Methodology: JJZ, ZHC, DMH, MZS, HMZ, CQL, JY. **Funding acquisition:** ZHQ, LS. **Supervision:** ZHQ. **Competing interests**: The authors declare that they have no competing interests. **Data and materials availability:** All data are available in the main text or the supplementary materials.

## Supplementary Materials

### Materials and Methods

#### Mice

C57BL/6J mice were obtained from Bestest (Zhuhai, China). The RCL-GCaMP6f mice (Ai95D; stock no. 024105; The Jackson Laboratory) were crossed with mice expressing Cre recombinase under the control of the EIIa promoter (stock no. 003724; The Jackson Laboratory) to achieve expression of GCaMP6f. Mice were mated overnight, and the morning when plugs were detected was considered embryonic day (E) 0.5. Mice were housed on a regular light–dark cycle and fed a standard diet with water provided ad libitum. All animal procedures were conducted in accordance with the guidelines of the Committee for the Use of Laboratory Animals, Guangdong Institute of Intelligence Science and Technology, China.

#### Cynomolgus macaques

Three pregnant cynomolgus macaques (weighing over 8.5 kg, and aged 9 years) were obtained from Guangdong Landau Biotechnology Co., Ltd. (Guangzhou, China) and housed in a conventional facility with one monkey per cage. The housing environment was maintained according to AAALAC (Association for Assessment and Accreditation of Laboratory Animal Care) guidelines, with an ambient temperature between 18°C and 26°C and relative humidity ranging from 60% to 80%. All housing and experimental procedures were approved by the Institutional Animal Care and Use Committee (IACUC) of the Guangdong Laboratory Primate Institute, adhering to ethical standards for animal welfare. The study was conducted under protocols approved by the Committee on the Ethics of Animal Experiments of the Guangdong Institute of Intelligence Science and Technology.

#### fUS imaging of embryonic heart and brain

Pregnant mice from E8.5 to E18.5 were used in this part of the experiment. Anesthetized pregnant mice were placed supine on a constant temperature insulation mat maintained at 37°C. Fur was removed from the region of the belly overlying the uterus, and a small vertical incision was made into the abdomen. Through this incision, embryos, still encased within their uterine sac, were extracted and placed outside the mouse’s abdominal cavity. To ensure that the heart and brain could be imaged on the same plane, the embryos were positioned with their dorsal sides facing up or down, which allowed for access to the sagittal plane with a probe. Embryos were covered with pure ultrasound gel without bubbles and imaged by an 18.5 MHz single crystal piezoelectric transducer (L22–14v, Vermon Inc., Tours, France). The frame rate of the acquisition speed was 20 Hz. To test the brain function response to auditory stimulation, while performing the fUS by plane wave, sound stimulation (1 kHz, 90 dB SPL; 10 kHz, 90 dB SPL; 16 kHz, 70 dB SPL) was applied every 110 seconds for 3 times.

Pregnant Cynomolgus macaques were slightly anesthetized by Zoletil@50. Fur was gently removed from the region of the belly overlying the uterus. The traditional B-mode and color doppler were first applied. After the location, the biparietal diameter (BPD) and the greatest length (GL) is measured, fUS is then applied. The circular wave imaging was obtained by using UD9800 system (Wuxi Hisky Medical Technologies Co., Ltd.) equipped with convex array (UDC5-1, central frequency = 3.5 MHz, R = 50 mm). 500 frames of circular wave imaging with 5 steering angles, from −10°to 10°, at the frame rate of about 8000 Hz. The plane wave imaging as obtained by using Verasonics system (Vantage 256, Verasonics Inc, Kirkland, USA) equipped with a 7.5 MHz single crystal piezoelectric transducer (L11–4v, Vermon Inc., Tours, France). While performing the fUS by plane wave, sound stimulation (1 kHz, 70 dB SPL) is applied every 110 seconds for 3 times. With a customed Arduino setup, the sound stimulus can be labeled correctly.

#### Calcium imaging of embryonic brain under sound stimulation

With a custom-built large-field calcium imaging platform capturing the calcium imaging from top, the Calcium imaging can be applied. Transgenic E18.5 mice was carefully exposed from its uterine sac while leaving the placenta intact. The embryo was placed on side so that the sagittal section of the mouse brain could be record. While performing the calcium imaging, sound stimulation (1 kHz, 90 dB SPL, 25 s duration; 10 kHz, 90 dB SPL, 25 s duration; 16 kHz, 25 s duration, 70 dB SPL) was applied every 110 seconds for 3 times.

#### Analysis

fUS data were analyzed using MATLAB by selecting regions of interest to extract blood flow signals from the frontal, central and posterior brain regions as well as the heart. Correlations between brain blood flow signals and between brain and cardiac blood flow signals were analyzed. A 0.3 Hz high-pass filter was applied to extract purer brain blood flow signals by removing low frequency noise. Deconvolution filtering was further utilized to remove interfering cardiac blood flow and harmonic components mixed within the brain signals.

Autoregressive power spectral estimation was performed over 10 s intervals on extracted frontal, central and posterior brain blood flow waveforms. Functional states for each 10 s interval were classified into three categories by comparing energy levels between 0.5-5 Hz and 6-10 Hz: focused in low frequencies (0.5-5 Hz), high frequencies (6-10 Hz), or without clear frequency features. These were respectively indicated by blue, purple and yellow stripes on the time-frequency map for long-term observation of changes in brain functional activity states. The proportions of the three functional states at different gestational ages were also statistically analyzed. Furthermore, the brain blood signals were converted into sound sequences according to the spectral distribution within every 10 seconds, in order to encode the time structure of the brain activity into sound sequences.

#### Behavioral test

P1 pups were obtained from C57BL/6J E18.5 pregnant mice purchased from Bestest (Zhuhai, China). The day before the experiment, the soundproof chamber was cleaned with alcohol and left open overnight to eliminate any odors. Prior to the experiment, the pups’ abdominal region was checked to ensure they were properly fed and capable of normal movement. A speaker for sound stimulation was set up inside the soundproof chamber, and the pups were placed inside the chamber. After allowing the pups to acclimate, sound stimuli at different frequencies (1 kHz, 90 dB SPL, 25 s duration; 10 kHz, 90 dB SPL, 25 s duration; 16 kHz, 25 s duration, 90 dB SPL) were administered, with each frequency presented once every 110 seconds for 3 times. Finally, the distance traveled by the P1 pups in response to the sound stimuli was measured and calculated.

#### Genotyping

GCaMP6f+/− heterozygous embryos were produced by intercrossing mutant RCL-GCaMP6f and wild-type parents. All genotyping were determined by PCR analysis of DNA extracted from embryonic tails. The following primers were used GCaMP6f Mutant Forward: ACG AGT CGG ATC TCC CTT TG; Wild type Forward: AAG GGA GCT GCA GTG GAG TA; Common: CCG AAA ATC TGT GGG AAG TC. PCR cycling conditions were as follows: initial denaturation at 94°C for 5 min, followed by 10 cycles of 94°C for 30 s, then 65°C (−1°C/cycle) for 30 s, 72°C for 50 s, followed by 25 cycles of 94°C for 30 s, 55°C for 30 s, 72°C for 50 s, followed by 72°C for 5 min, and then a final hold at 16°C. Reactions were separated on 2% agarose gels, yielding the following band sizes: WT, 297 bp; Mutant = ∼450 bp; Heterozygote = ∼450 bp and 297 bp.

#### Immunohistochemical Fluorescent Staining

For E8.5 embryos, following dissection from the uteri of pregnant mice, post-fixation was performed at room temperature for 4 hours using 4% PFA. Following post-fixation, specimens were rinsed with PBS. Embryos were then incubated overnight at 4°C in a blocking solution containing 10% normal goat serum, 1% bovine serum albumin, and 0.3% Triton X-100. After blocking, embryos were incubated with CD31 antibody (1:50; Cat. No. ab7388, Abcam) for two nights at 4°C and then with Alexa Fluor 555-conjugated Donkey anti-rat secondary antibody (1:50; Cat. No. A11077, Invitrogen). Light-sheet images were captured by LiTone XL Light-sheet Microscope (Light Innovation Technology Limited) with a 10X objective (N.A.= 0.6). Further image processing was performed by LitScan 3.2.0 (Light Innovation Technology Limited).

---

**Fig. S1.**
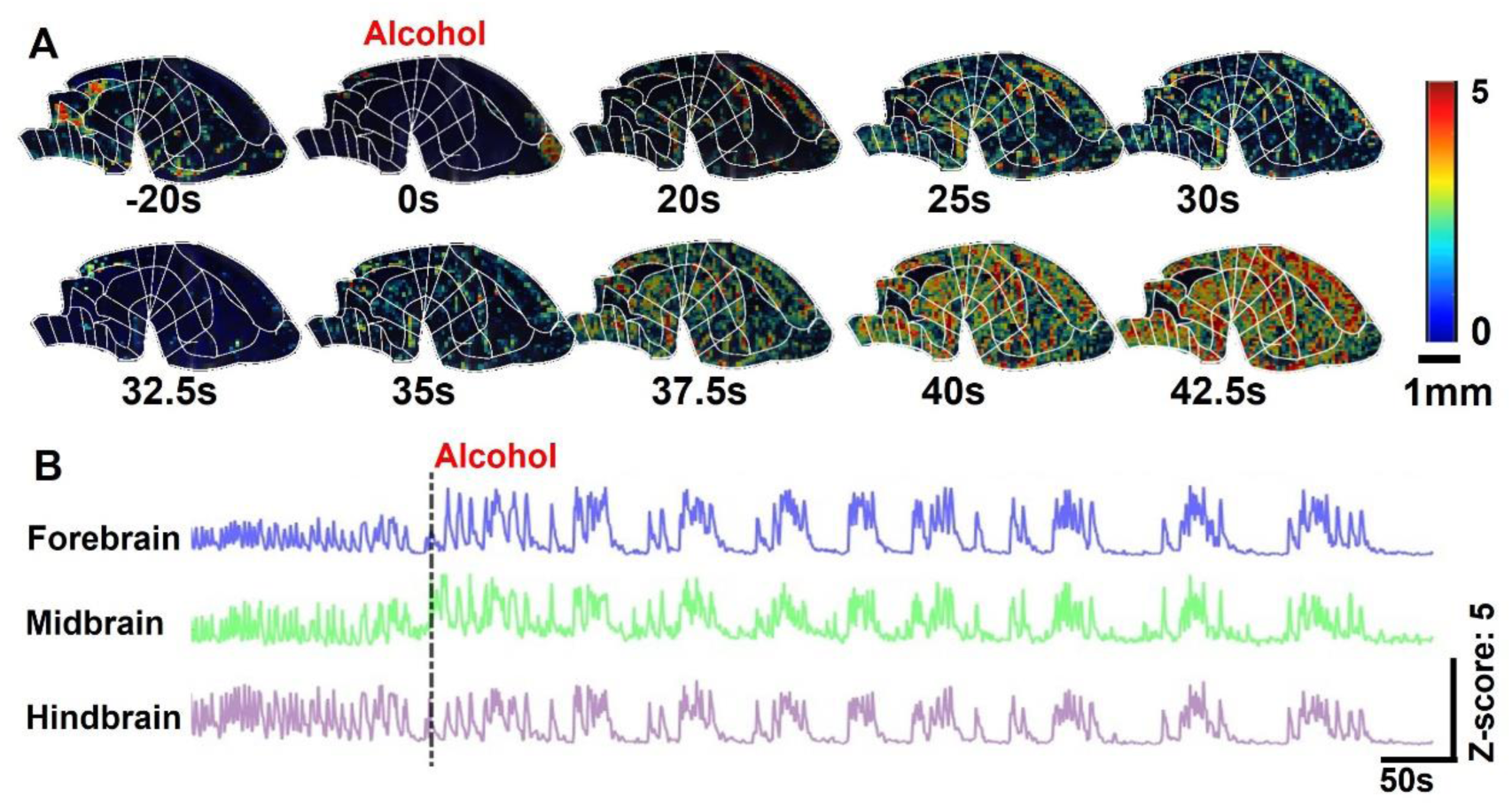
Embryonic mouse brain strongly response to alcohol administration in pregnant mouse. (A) Representative fUS images showing an embryo brain’s response to alcohol (3 g/kg) administration in a pregnant mouse. Scale bar: 1mm. (B) Z-score trace of fUS signals from the forebrain, midbrain, and hindbrain (top to bottom) following alcohol administration (3 g/kg) in pregnant mice.

**Fig. S2.**
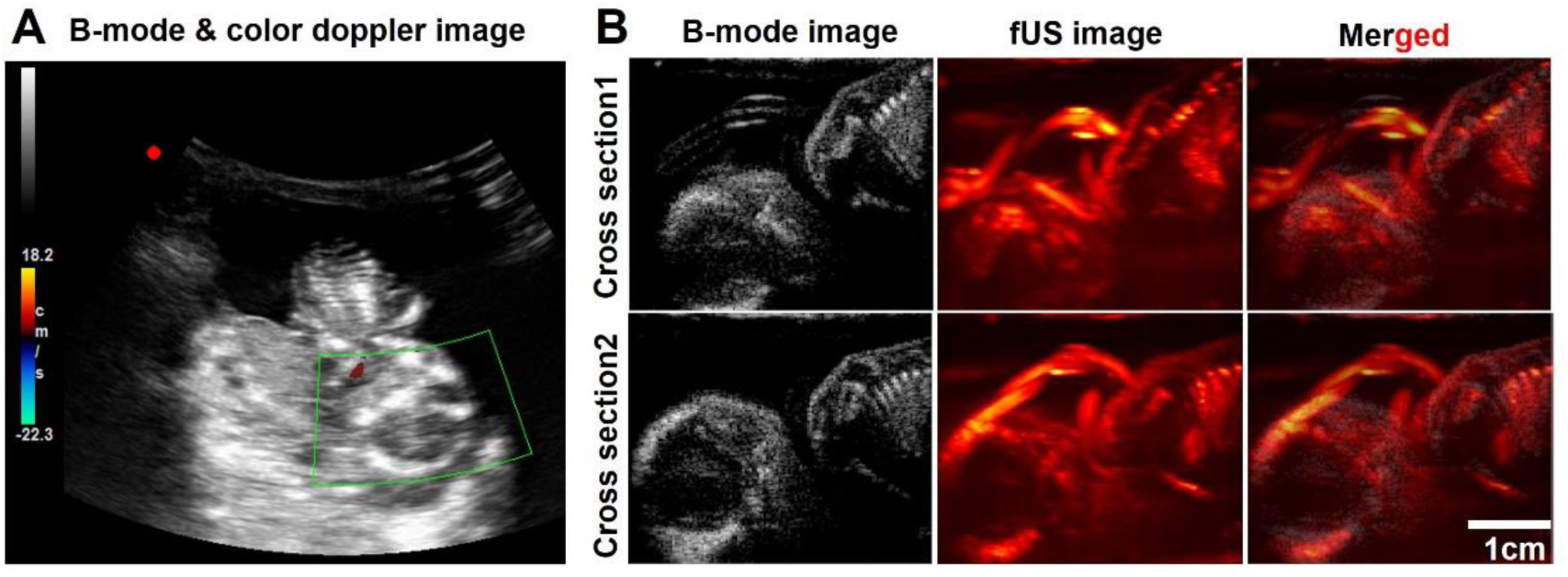
B-mode (3.5 MHz) and fUS (7.5 MHz) imaging of a cynomolgus macaque embryo at gestation day 30. (A) Traditional B-mode and Doppler imaging using a line-by-line scanning method. (B) Plane waves compounded B-mode imaging and fUS imaging of the embryo. The B-mode images reveal the major body structures, while the fUS provides more detailed visualization of brain structures and clearly shows umbilical cord entanglement. These images demonstrate the advantages of fUS for detailed embryonic monitoring.

